# Small-molecule sensitization of RecBCD helicase-nuclease to a Chi hotspot-activated state

**DOI:** 10.1101/2020.05.04.077768

**Authors:** Ahmet C. Karabulut, Ryan T. Cirz, Gerald R. Smith

## Abstract

Coordination of multiple activities of complex enzymes is critical for life, including transcribing, replicating, and repairing DNA. Bacterial RecBCD helicase-nuclease must coordinate DNA unwinding and cutting to repair broken DNA. Starting at a DNA end, RecBCD unwinds DNA with its fast RecD helicase on the 5’-ended strand and its slower RecB helicase on the 3’-ended strand. At Chi hotspots (5’GCTGGTGG3’), RecB’s nuclease cuts the 3’-ended strand and loads RecA strand-exchange protein onto it. We report here that a small molecule NSAC1003, a sulfanyltriazolobenzimidazole, mimics Chi sites by sensitizing RecBCD to cut DNA at a Chi-independent position a certain percent of the DNA substrate’s length. This percent decreases with increasing NSAC1003 concentration. Our data indicate that NSAC1003 slows RecB and sensitizes it to cut DNA when the leading helicase RecD stops at the DNA distal end. Two previously described RecBCD mutants altered in the RecB ATP-binding site also have this property, but uninhibited wild-type RecBCD lacks it. Computation docks NSAC1003 into the ATP-binding site, suggesting that NSAC1003 acts directly on RecB. NSAC1003 will help elucidate the molecular mechanisms of RecBCD-Chi regulation and DNA repair. Similar studies could help elucidate other DNA enzymes whose activities are coordinated at chromosomal sites.

## INTRODUCTION

Complex, multi-subunit enzymes are required for most activities on nucleic acids, including replication, recombination, DNA repair, transcription, RNA splicing, and protein synthesis. Particularly important is the repair of DNA double-strand breaks (DSBs) because when their DNA is broken, cells must repair it or they die. Faithful DSB repair requires DNA helicases and nucleases to prepare the broken DNA for interaction with an intact homologous DNA molecule. If the interacting DNA molecules differ genetically, DSB repair can also produce genetic recombinants and thereby propel evolution. Understanding the molecular basis of DSB repair and recombination requires understanding how complex, multi-functional enzymes act on DNA. Here, we describe a small organic molecule that alters the activity of a DNA helicase-nuclease complex in a novel way that lends new insights into the enzyme’s mechanism and regulation.

In *Escherichia coli* and other enteric bacteria, DSB repair and recombination require RecBCD, a complex three-subunit 330 kDa enzyme with both DNA helicase and DNA nuclease activities (1–3). Its multiple activities are involved in the initial steps of DSB repair and recombination (Figure 1A). RecBCD binds tightly to a ds DNA end and unwinds it rapidly (up to ~1.5 kb/sec) and with high processivity (up to 100 kb without dissociating) (4). RecB helicase binds to the 3’ end, and RecD to the 5’ end (5,6). Each helicase subunit hydrolyses ATP and moves along its bound strand (7). Because RecB is slower than RecD, a single-stranded (ss) DNA loop accumulates, presumably ahead of the RecB subunit (4). This loop grows with time of incubation, as do the short and long ss tails extending behind the enzyme. Annealing of the tails forms ds DNA and a second loop; the two loops continue to grow and move with time of incubation. Upon encountering a Chi site (5’ GCTGGTGG 3’), a hotspot of recombination, the RecB nuclease cuts the strand with this sequence (8). Unwinding continues, and RecA strand-exchange protein is loaded onto the newly generated 3’-ended strand likely by the RecB nuclease domain (9,10). The RecA-ssDNA filament invades intact homologous DNA to form a displacement (D)-loop (11). Subsequent reactions involve formation and resolution of a Holliday junction into reciprocal recombinants or priming of DNA synthesis to form a non-reciprocal recombinant (12,13).

**Figure 1.**
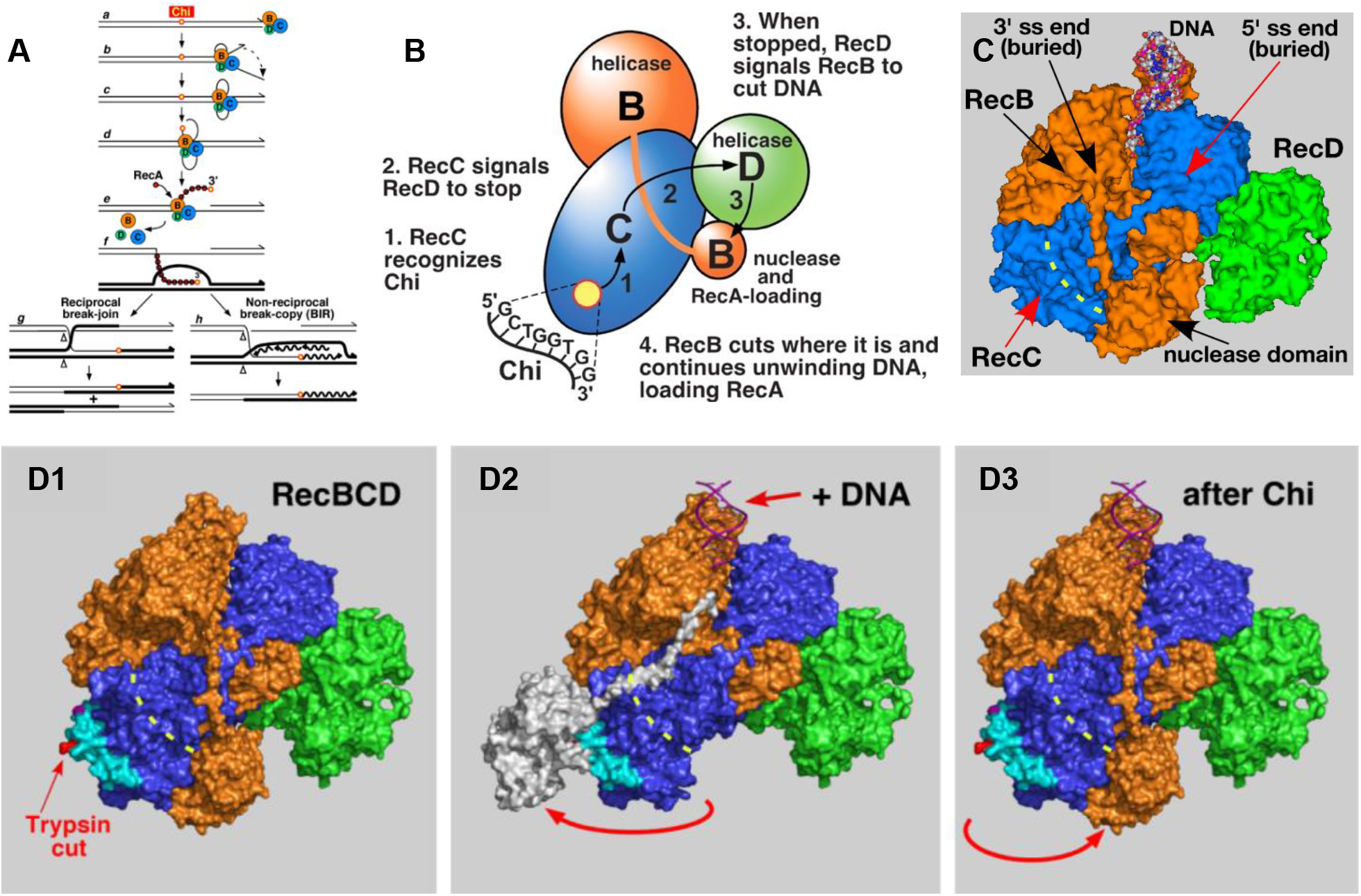
Models for RecBCD enzyme and its Chi-dependent promotion of DNA break repair and genetic recombination. **(A)** RecBCD pathway of genetic recombination. See Introduction for explanation. **(B)** Signal-transduction model for Chi’s control of RecBCD enzyme (14). When Chi is in the RecC tunnel, RecC signals RecD to stop unwinding DNA; RecD signals RecB to nick the DNA and to begin loading RecA. **(C)** Crystal structure of RecBCD bound to a ds DNA hairpin (PDB 1W36; (6)). RecB (orange) contains helicase and nuclease domains connected by a tether. RecD (green) is held to RecB via RecC (blue). During unwinding, the 3’-ended strand passes through a tunnel (yellow dashed line) in RecC and into the nuclease active site when Chi is encountered. **(D)** Nuclease-swing model for Chi’s control of RecBCD enzyme (15). Before DNA is bound (D1), RecBCD in solution assumes its conformation in the published structures (PDB 1W36 and PDB 5LD2). Upon binding DNA (D2), the nuclease domain swings away from the exit of the RecC tunnel (15). When Chi is encountered during unwinding (D3), the nuclease domain swings back, cuts the DNA at Chi, and begins loading RecA protein, perhaps after rotating to prevent further nuclease action. Modified from (16).

The molecular mechanism by which a Chi hotspot signals RecBCD to cut DNA at Chi and to begin loading RecA remains to be fully elucidated. Based on the behavior of two mutants altered in the RecB helicase domain, Amundsen et al. (14) proposed a “signal-transduction” model (Figure 1B). In this model when Chi is in a tunnel in RecC (Figure 1C), RecC signals RecD to stop; RecD then signals RecB to cut the DNA at Chi and to load RecA. Subsequent analysis suggested that upon receiving the signal from RecD, RecB’s nuclease domain swings on its 19-amino-acid tether from an inactive position (on the “left” side of RecC) to its nuclease-active position (at the exit of the tunnel in RecC) (Figure 1D) (15,16). Extensive mutational analyses of the RecC tunnel and of the RecB tether support this model (17,18,16). Mutants with altered contacts between RecC-RecD, RecD-RecB, and RecB-RecC indicate that each of these contacts is important for Chi hotspot activity (19).

Here, we report the action of a small organic molecule on RecBCD, which mimics the behavior of the two RecB helicase mutants noted above (14). This small molecule, NSAC1003 (MW = 431; Figure 2A), was discovered by Achaogen in a screen for inhibitors of the helicase-nuclease activity of RecBCD. In the current studies we noted that it induces RecBCD to cut DNA at a novel position that depends on the NSAC1003 concentration and on the length of the substrate. The two RecB helicase mutants are altered in single amino acids (Y803H and V804E) near RecB’s ATP-binding site in a cryoEM structure (20) (see Discussion and Figure 5). These two helicase mutants also cut DNA at novel positions that depend on the substrate length (14). From these and other data, we infer that NSAC1003 has two effects: it slows the RecB helicase in a concentration-dependent manner, and it sensitizes the enzyme to cut DNA when RecD stops unwinding DNA at the end of the substrate. We discuss a molecular basis for these effects and how related compounds may also act on RecBCD and other complex enzymes.

**Figure 2.**
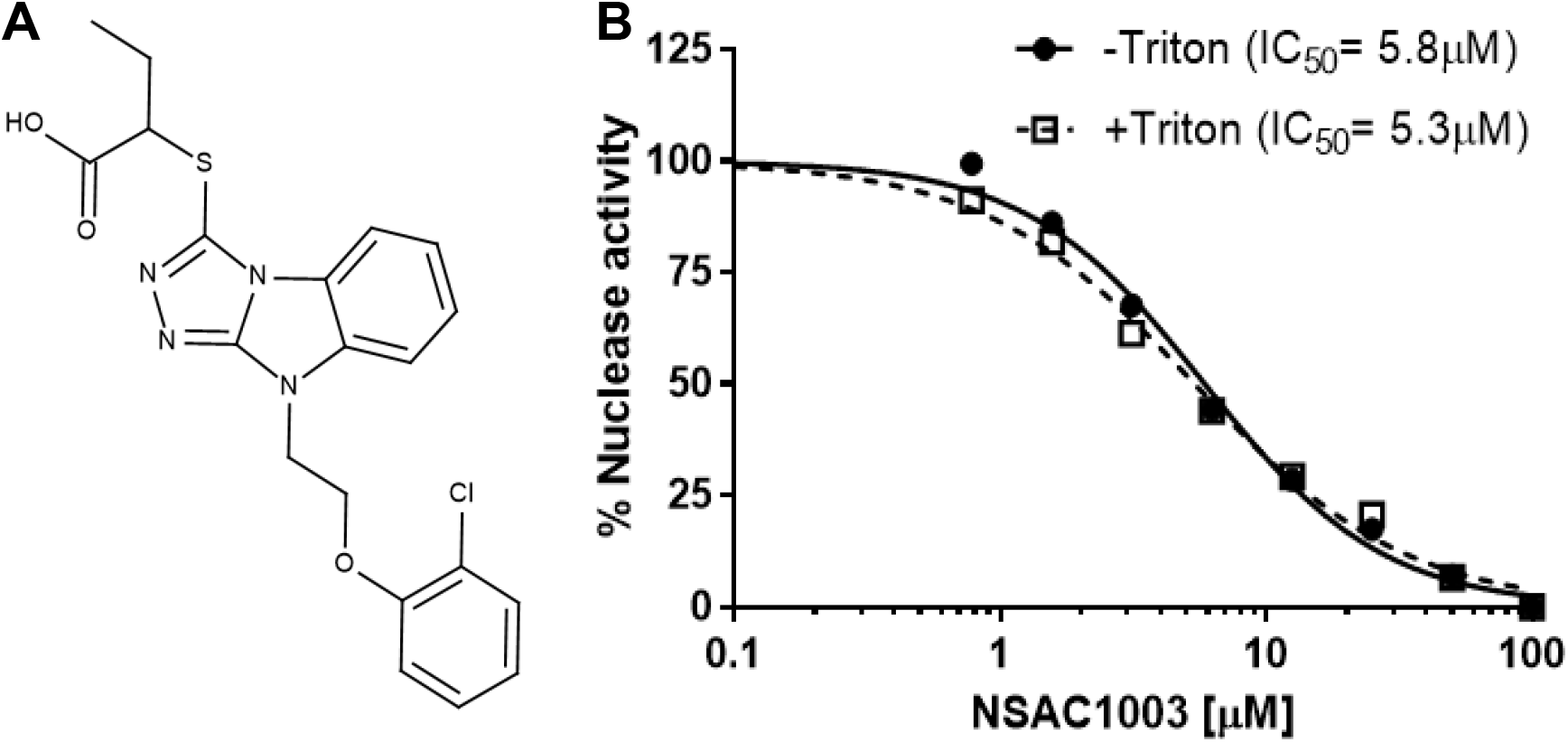
NSAC1003 inhibits RecBCD Chi-independent nuclease activity. **(A)** Structure of NSAC1003, a sulfanyltriazolobenzimidazole. The IUPAC name is 2-({7-[2-(2-chlorophenoxy)ethyl]-2,4,5,7-tetraazatricyclo[6.4.0.0^2^,^6^]dodeca-1(8),3,5,9,11-pentaen-3-yl}sulfanyl)butanoic acid. **(B)** RecBCD nuclease activity was assayed, using [^3^H] phage T7 DNA, in the presence of the indicated concentration of NSAC1003 with and without Triton X-100 (0.01%).

## MATERIALS AND METHODS

### Materials

RecBCD enzyme (7), pBR322 (*χ*° and *χ*^+^*E224*) (21), and [^3^H] phage T7 DNA were generous gifts from A.F. Taylor. Restriction enzymes and polynucleotide kinase were purchased from New England BioLabs. [γ-^32^P] ATP was purchased from Perkin-Elmer. Triton X-100 was purchased from Sigma. NSAC1003 was dissolved in DMSO at 10 mM; pure DMSO was added to experimental reactions to make a total concentration of 5% (nuclease activity in Figure 2) or 2% (unwinding and DNA cutting in Figure 3).

**Figure 3.**
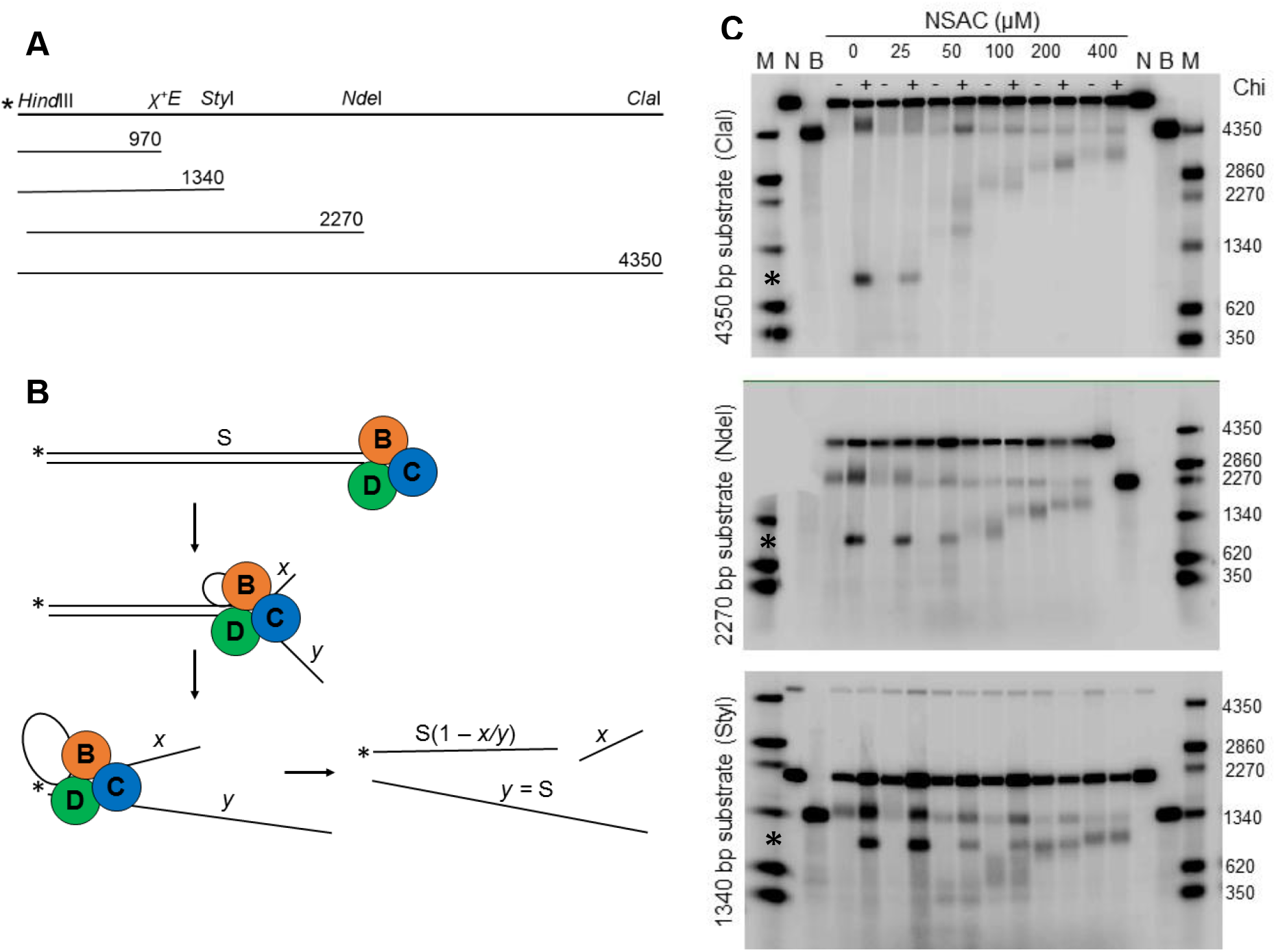
NSAC1003 induces Chi-independent DNA cuts. (**A**) Diagram of 5’-labeled (**\***) ds DNA substrates of various lengths. **(B)** Diagram of RecBCD unwinding DNA with production of a ss DNA loop and two tails (4). *x* is the rate of the RecB helicase on the 3’-ended (top) strand, and *y* the rate of the faster RecD helicase on the 5’-ended (bottom) strand. When RecD reaches the left end of the DNA, in the presence of NSAC1003 RecB makes a novel cut where it is at that moment, or *x/y* of the substrate length, measured from RecBCD’s entry point. The length of the labeled product [S(1 – *x/y*)] is a nearly constant fraction independent of the initial substrate length S. NSAC1003 inhibits RecB helicase more than RecD helicase; thus, *x/y* decreases as [NSAC1003] increases and the cut product is longer. **(C)** pBR322 DNA (*χ*° or *χ*^+^*E224*) was cut with *Hin*dIII and labeled at the 5’ end; portions were cut with *Cla*I, *Nde*I, or *Sty*I to produce singly labeled DNA ~4350, 2270, or 1340 bp long (panel A). DNA was reacted with RecBCD enzyme with the indicated concentrations of NSAC1003 for 60 sec. Reaction products were examined by gel electrophoresis as described in STAR methods. M, ss DNA markers; N, native (ds) substrate; B, boiled (ss) substrate; **\***, Chi-dependent product; -, *χ*° substrate; +, *χ*^+^*E224* substrate. The upper left corner of the middle gel was marred, obscuring some marker bands.

### Enzymatic methods

RecBCD “general” (Chi-independent) nuclease activity (Figure 2B) was assayed as described (22). Reaction mixtures (50 μL) contained 50 mM Tris-HCl (pH 8.5), 10 mM MgCl_2_, polyvinylpyrrolidone (1 mg/ml), 1 mM dithiothreitol, 25 μM ATP, RecBCD enzyme (4 units/ml), [^3^H] phage T7 DNA (2 μg/ml), and NSAC1003 at the indicated concentration; DMSO, the solvent for NSAC1003, was added to 5% final concentration in all reactions. Where indicated, Triton X-100 was added to 0.01%. Reactions were assembled on ice and transferred to a 37 °C bath to initiate the reaction. After 20 min at 37 °C, the reaction was stopped by addition of 25 μL of calf thymus DNA (0.2 mg/ml) and 250 μL of 5% trichloroacetic acid (TCA). After 10 min on ice, the tubes were centrifuged for 5 min at 13,400 RPM, and 300 μL of the supernate were assayed with a scintillation counter.

Chi-cutting activity (Figure 3C) was assayed as described (14). Reaction mixtures (15 μL) contained 25 mM Tris-HCl (pH 7.5), 2.5 mM MgCl_2_, 1 mM dithiothreitol, 0.2 nM [5’-^32^P] DNA, 0.4 nM RecBCD enzyme, and NSAC1003 at the indicated concentration. DMSO was present at 2% in all reactions, which were incubated at 37 °C for 10 min before initiating the reaction by addition of ATP to 5 mM. After 60 sec at 37 °C, the reaction was stopped by addition of 5 μL of stop buffer, and the products analyzed by electrophoresis on a 1.5% agarose gel in TAE buffer (110 V for 3 hr). The gel contents were analyzed by Typhoon Trio PhosphorImager (GE Lifesciences) and ImageQuant TL software (GE Lifesciences).

### Docking methods

Using DockingServer (23), NSAC1003 was docked to a 20 Å cube containing the RecB ATP site in cryoEM structure PDB 5LD2 (24) with ADPNP and Mg^2+^ removed. Gasteiger partial charges were added to the ligand atoms. Non-polar hydrogen atoms were merged, and rotatable bonds were defined. Essential hydrogen atoms, Kollman united atom-type charges, and solvation parameters were added with the aid of AutoDock tools (25). Affinity (grid) maps of 20 Å grid points and 0.375 Å spacing were generated using the Autogrid program (25). During the search, a translational step of 0.2 Å and quaternion and torsion steps of 5 were applied. AutoDock parameter set- and distance-dependent dielectric functions were used in the calculation of van der Waals and electrostatic terms, respectively. Docking simulations were performed using the Lamarckian genetic algorithm (LGA) and a local search method (26). Initial position, orientation, and torsions of the ligand molecules were set randomly. All rotatable torsions were released during docking. Each docking experiment was derived from 100 different runs set to terminate after a maximum of 2,500,000 energy evaluations. The population size was set to 150. The docking with the highest affinity score is shown in Figure 5.

## RESULTS

### NSAC1003 inhibits RecBCD nuclease activity

RecBCD has potent Chi-independent (“general”) nuclease activity under conditions with excess Mg^2+^ relative to ATP (22). In standard assays of RecBCD nuclease activity with 10 mM Mg^2+^ and 25 μM ATP, we found that NSAC1003 inhibited with an IC_50_ of 5.8 μM (Figure 2B). Because some organic compounds found in screens for antibiotics form microcrystals or aggregates and may sequester an enzyme rather than simply inhibit it (27), we repeated the assays in the presence of 0.01% Triton X-100, which is thought to counter the effect of microcrystals. NSAC1003 inhibition was indistinguishable in the presence and absence of Triton. Furthermore, addition of NSAC1003 (5 – 80 μM) to RecBCD, either without DNA or with DNA in an active reaction, followed by centrifugation (14,000 ×*g*, 15 min) did not remove RecBCD from solution (assayed by Western blots for RecB and RecC; A.F. Taylor, pers. comm.). A compound closely related to NSAC1003 inhibits the RecBCD helicase-nuclease activity in an ATP-competitive manner (R. Cirz, unpublished data), suggesting that these compounds inhibit ATP hydrolysis, which is required for helicase activity and thus nuclease activity (22,28,29,7). Thus, we conclude that NSAC1003 inhibits RecBCD by direct binding, perhaps to the ATP-binding site(s) (see Discussion and Figure 5).

### NSAC1003 induces RecBCD to cut DNA at novel positions

Under conditions with excess ATP relative to Mg^2+^, RecBCD unwinds DNA and nicks one strand near Chi to generate a hotspot of recombinational exchange at Chi, as in cells (1). Using 5 mM ATP and 2.5 mM Mg^2+^, we tested the effect of NSAC1003 on RecBCD’s DNA unwinding and Chi-cutting activities. The linear ds DNA substrate was labeled at one 5’-end with ^32^P and contained, or not, an internal Chi site (*χ*^+^*E224*) (Figure 3A). After brief reaction, the products were analyzed by gel electrophoresis in comparison with ds and ss DNA length standards. In the absence of NSAC1003 RecBCD unwound some of the Chi-containing DNA (4.35 kb long; top panel) and produced a radioactive ss DNA fragment ~970 nucleotides long, as expected from the Chi site being ~970 bp from the 5’-^32^P label; also as expected, this fragment was not detected with DNA substrate lacking Chi (Figure 3B, C). (RecBCD entering the DNA from the right but not from the left, as drawn, cuts at Chi (30).) Similar results were found with 25 μM NSAC1003. With 50 – 400 μM NSAC1003, however, a Chi-dependent fragment was not detected but was replaced with a longer fragment indicative of cutting before Chi. This fragment’s length increased with increasing NSAC1003 concentration and was observed with and without Chi. We interpret these results below.

With shorter DNA substrates, the results changed in an interesting way. With the 2.27 kb substrate (middle panel), the Chi-dependent fragment (~970 nucleotides long) was detected with 0, 25, and 50 μM NSAC1003 but not with 100, 200, or 400 μM NSAC1003. At these higher NSAC1003 concentrations, radioactive fragments of increasing length were produced with DNA containing Chi or not. The length of these fragments increased with increasing NSAC1003 concentration, as noted above with the 4.35 kb substrate. With the 1.34 kb substrate (bottom panel), the Chi-dependent fragment was detected with 0 to 100 μM NSAC1003. An additional, Chi-independent fragment was produced with 50 to 400 μM NSAC1003, and as above its length increased with increasing NSAC1003 concentration.

### The positions of NSAC1003-induced cuts depend on the NSAC1003 concentration and on the substrate length

The lengths of the Chi-independent fragments noted above depended on the NSAC1003 concentration in a Michaelis-Menten sort of way, with an apparent half-maximal effect at ~50 – 100 μM (Figure 4A). This value is higher than the IC_50_ for NSAC1003 inhibition of the “general” (Chi-independent) nuclease activity (5.8 μM; Figure 2), likely because ATP competes with NSAC1003; the ATP concentration was 25 μM for the nuclease assay and 5 mM for the unwinding and cutting assay.

**Figure 4.**
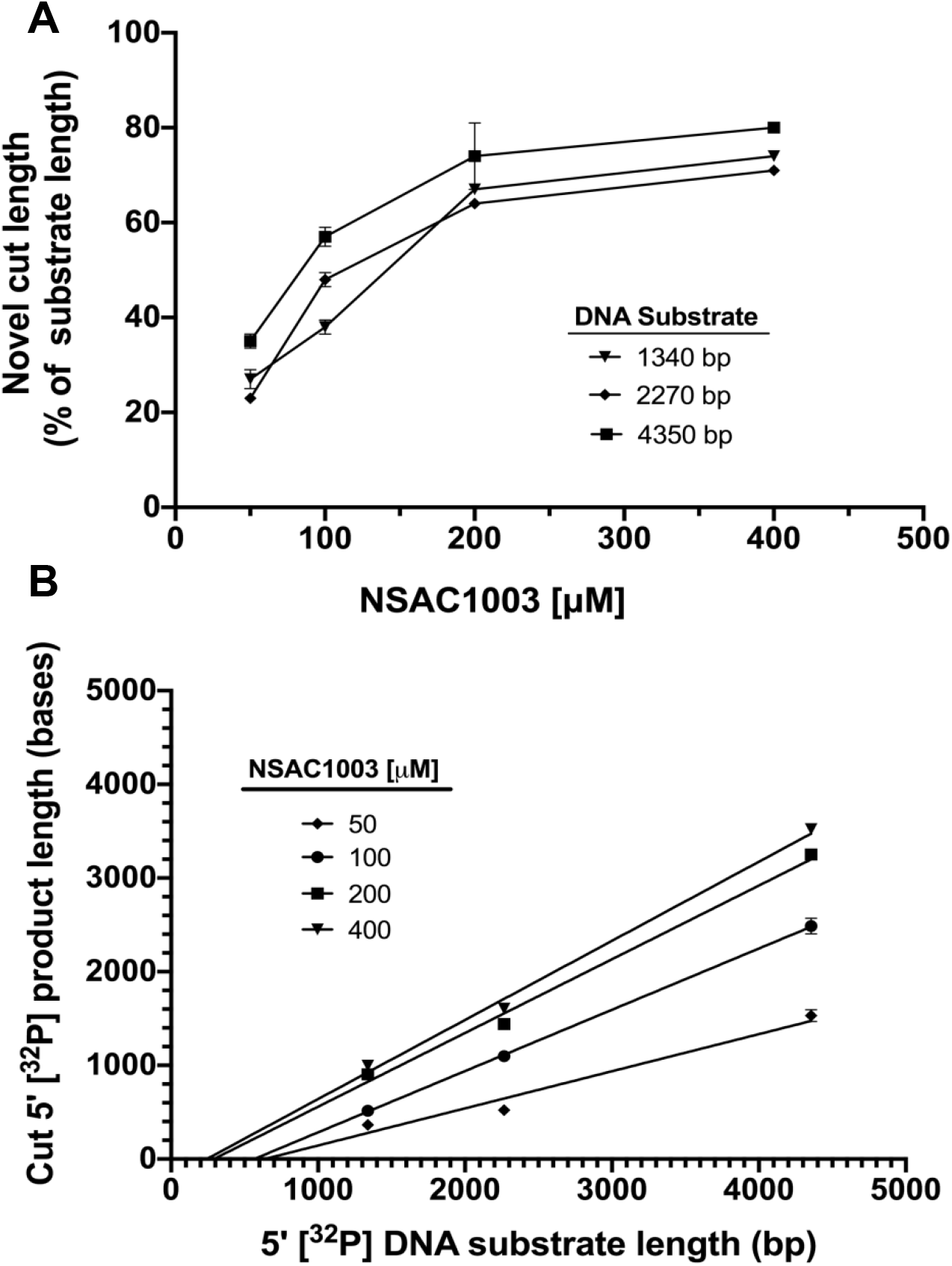
The length of the NSAC1003-induced cut product is a linear function of substrate length, dependent on the NSAC1003 concentration. Three experiments of the type shown in Figure 3 were analyzed; data are the mean ± SEM (not visible in most cases). The length of the cut product (middle of the band) was estimated by interpolation against the indicated size markers (boiled substrates and products of digestion with *Sal*I, *Bam*HI, or *Alw*NI). **(A)** Labeled product length increases with increasing NSAC1003 concentration. **(B)** Labeled product length is a linear function of substrate length. Note that the extrapolate to zero substrate length is slightly negative (−250 bases at 50 μM; −370 at 100 μM; −290 at 200 μM; and −200 at 400 μM) (see Discussion).

The radioactive product lengths were a linear function of the length of the DNA substrate (Figures 3B and 4B). These data show that NSAC1003 induces RecBCD to cut the DNA ~30 – 70% of the distance from the entry point (the unlabeled end of the substrate) (Figure 3). In other words, at a low concentration (25 μM) of NSAC1003 the enzyme cuts at ~70% of the substrate length from the entry point, and at higher concentration it cuts at ~30% of the substrate length. This outcome is nearly the same for each substrate length tested. These results suggest that NSAC1003 slows RecB helicase, relative to RecD, and induces RecB nuclease to cut the DNA where it is when RecD reaches the end of the DNA (see Figure 3B and Discussion).

## DISCUSSION

Our results presented here show a remarkable similarity between the effect of NSAC1003 inhibitor and the effect of two mutations altering amino acids in the RecB helicase very near its ATP-binding site (Figure 5). Molecular docking (23) indicates that NSAC1003 binds to the ATP-binding site in RecB (Figure 5) with high affinity (0.3 μM) and with two consequences: the RecB helicase is slowed, likely reflecting the competition with ATP binding and hydrolysis, and the nuclease is sensitized to cut the DNA when RecD helicase stops, perhaps because NSAC1003 alters the RecBCD conformation in the same way as the two mutations in RecB (Y803H and V804E) (14). These effects of NSAC1003 indicate that this compound will be informative in further elucidating Chi’s control of RecBCD enzyme and other complex, multi-activity enzymes, as discussed below.

**Figure 5.**
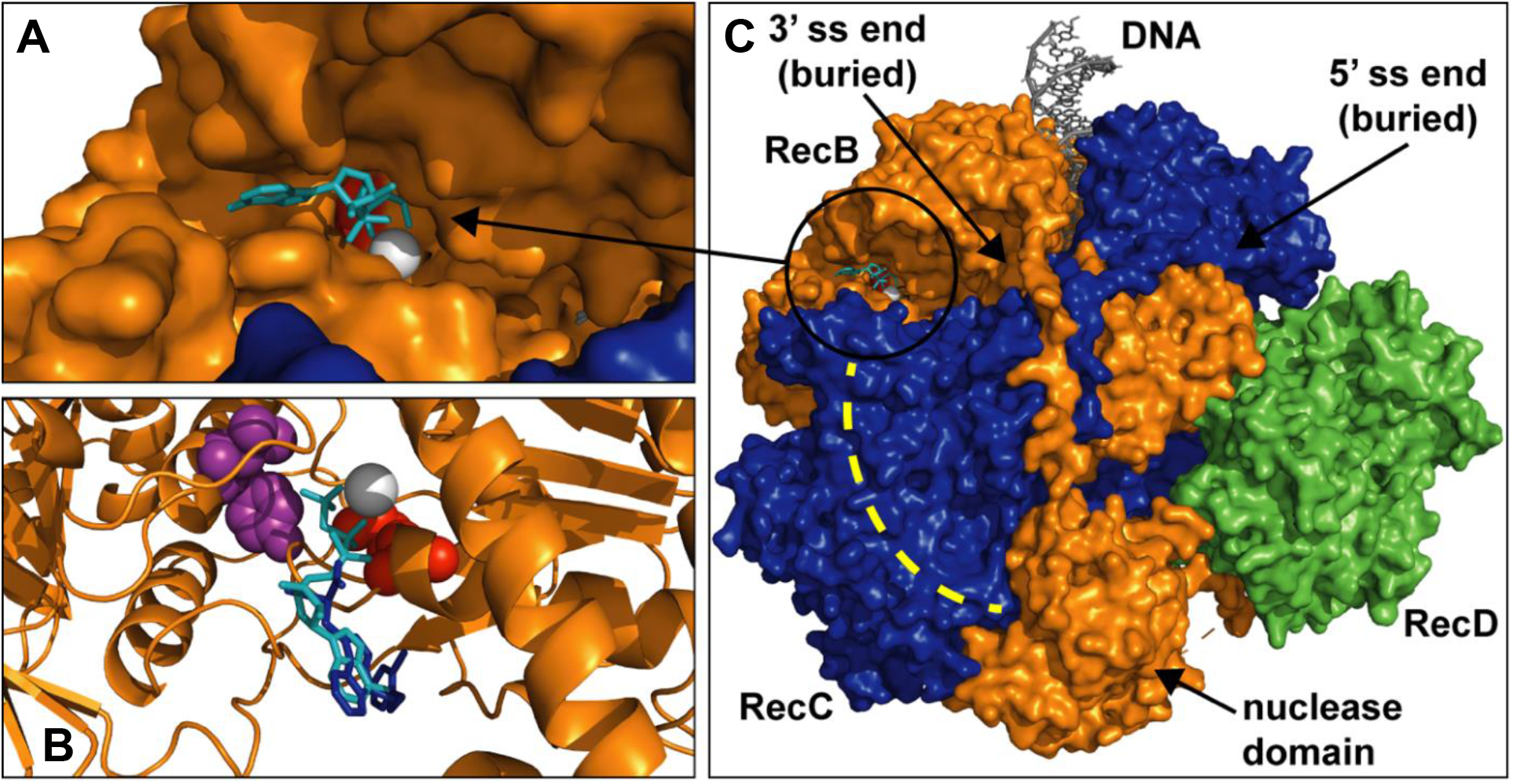
RecB helicase mutants and NSAC1003 docking site are at the RecB ATP-binding site. **(A)** Part of the structure of RecB (orange, surface view from PDB 5LD2) is shown with the ATP analog adenosine 5′-[β,γ-imido]triphosphate (ADPNP) (cyan sticks) and the catalytic Mg^2+^ (grey sphere). **(B)** As in panel A with NSAC1003 (dark blue sticks) docked with the Docking Server (23) to a 20 Å cube of RecB centered on the ATP-binding site but without ADPNP or Mg^2+^ for the calculation and superimposed with ADPNP structure (cyan). Walker A box lysine K29 (red spheres) and two helicase mutant positions (Y803 and V804, purple spheres) are also shown. **(C)** Structure of RecBCD (RecB, orange; RecC, blue; RecD, green; surface view of PDB 5LD2 with ADPNP) showing the position of the RecB ATP-binding site (circled) and the RecC tunnel (dashed yellow line) in which Chi is recognized.

Our interpretations of the data in Figures 3 and 4 reflect those of the two RecB helicase mutants that cut DNA at a certain fraction of the length of the DNA substrate (14). Initially, it was mysterious how these mutants could measure the length of the substrate, calculate a certain percent of that length (~19% for Y803H and ~6% V804E), and cut the DNA at that position. Analysis of the rates of the mutant RecB and RecD helicases by electron microscopy of partially unwound DNA molecules showed that the ratio of the RecB:RecD helicase rates was nearly the same as the fraction of the substrate length (from the RecBCD entry point) at which cuts occur. This led to the conclusion that, when RecD stops at the end of the DNA, it signals RecB to cut where it is at that moment. We propose the same interpretation for the effect of NSAC1003 – it slows RecB in a concentration-dependent manner and sensitizes RecB to cut the DNA when RecD stops at the end of the DNA. The high similarity in the effects of the mutations and of NSAC1003 is consistent with the altered amino acids being very near the ATP binding site in RecB and, we infer, NSAC1003 binding to the ATP site in a competitive manner (Figure 5). Slowing of the RecB helicase by the mutations and by NSAC1003 is thereby readily accounted for.

The mechanism for sensitization of RecB to cut the DNA when RecD stops is less obvious. Recent analysis of mutations in each subunit that reduce or block Chi hotspot activity shows that amino acids widely scattered throughout the large RecBCD complex are important for Chi to signal DNA cutting (19). Three points of contact (RecC-RecD, RecD-RecB, and RecB-RecC) appear to act sequentially to transmit the Chi signal from the RecC tunnel to the RecB nuclease domain (Figure 1B, 1C, and 1D). NSAC1003 and the RecB helicase mutants could bypass the early steps of this cascade (Chi recognition in the RecC tunnel, and Chi-bound RecC signaling RecD to stop) and directly sensitize RecB to cut the DNA when RecD stops in the same way that Chi signals wild-type RecD, when stopped, to induce RecB to cut where it is at that moment [~4 – 6 nucleotides to the 3’ side of the Chi sequence bound in the RecC tunnel (Figures 1 and 3) (30).

Two features of the cut products lead to further insights into the control of RecBCD nuclease activity. NSAC1003-induced cutting, like Chi-independent cutting by the RecB helicase mutants (Y803H and V804E), produces a smear of DNA fragments differing in length by ~200 – 300 nucleotides (Figure 3C) (14). Chi-dependent cutting, however, occurs over only a 2- to 3-nucleotide range, ~4 – 6 nucleotides 3’ of the Chi sequence (30). We infer that the signal from Chi acts much more quickly than the signal from RecD stopping at the end of DNA (by NSAC1003 or by the RecB helicase mutants). Specifically, we propose that the time it takes for the RecB nuclease domain to swing from its position on the “left” side of RecBCD to the DNA exit of the tunnel in RecC (Figure 1D) is ~1 msec after Chi’s encounter but ~200 msec after RecD stops in the RecB helicase mutants or in the presence of NSAC1003. These estimates follow from RecBCD’s unwinding rate of ~1 bp/msec (4). Upon encountering Chi, the nuclease would swing to the tunnel exit in ~1 msec, with variability from molecule to molecule to account for the spread of Chi-dependent cuts over ~3 nucleotides. With NSAC1003 or the RecB helicase mutants, the swing would take much longer (~200 – 300 msec) and allow RecB to advance ~200 – 300 nucleotides before cutting. This interpretation accounts for the Chi-dependent band being much sharper than the NSAC1003- or RecB helicase mutant-dependent bands (Figure 3) (14). An alternative interpretation is that Chi may form a kinked configuration in the RecC tunnel (20) and slow or stop DNA translocation until the DNA is cut near Chi; if so, a longer nuclease swing-time would still result in cuts only a few nucleotides from the Chi sequence. In experiments without Chi, the DNA in the RecC tunnel would not typically be kinked when RecD reaches the DNA end, and NSAC1003- or helicase mutant-induced cuts would be spread over a larger region from the same nuclease swing-time. Whether Chi is kinked in the RecC tunnel at the moment of DNA cutting in an active RecBCD reaction has not been reported.

The second feature of the Chi-independent cuts is the non-zero extrapolate of the position of the cuts. For both NSAC1003 and the RecB helicase mutants, the extrapolate to zero substrate length is ~ −200 – −300 nucleotides (Figure 4B) (14). Thus, the cuts occur ~200 – 300 nucleotides farther along the DNA than expected from the cuts being exactly a constant fraction of the substrate length. This distance is that expected from the rather slow movement of the nuclease domain inferred from the smear distribution noted above. Both features concur in predicting that RecB advances, after RecD stops at the DNA end, ~200 – 300 nucleotides before cutting in the presence of NSAC1003 or in the RecB helicase mutants.

These features of NSAC1003 action on RecBCD should enable further biophysical analysis of RecBCD and AddAB helicase-nucleases, closely related DNA repair enzymes ubiquitous among bacteria but not reported in eukaryotes (31). In particular, the predicted relatively slow speed of swinging of the RecB nuclease domain should be easier to detect with NSAC1003 or the RecB helicase mutants than with Chi and wild-type RecBCD. RecBCD is a member of the superfamily 1 helicases, which, like many ATPases, have well-conserved motifs at their ATP-binding sites (32,33). RecB and RecD have a so-called Walker A box, which binds to ATP (Figure 5); mutations in each Walker A box, such as RecB K29Q (Figure 5) show that both are required for ATPase and helicase activity (28,29,7). Our inference that NSAC1003 binds to the RecB ATP-binding site (Figure 5) therefore predicts that NSAC1003 would inhibit other helicases. If so, this compound may be broadly useful in studying ATP-hydrolyzing molecular motors and ATPases in general, one of the largest classes of enzymes, including DNA and RNA polymerases.

## ACKNOWLDEGMENTS

We are grateful to Andrew Taylor and Sue Amundsen for purified RecBCD enzyme, DNAs, and advice on their use; to Andrew Taylor for unpublished data; and to Sue Amundsen, Randy Hyppa, Brett Kaiser, and Rasi Subramaniam for helpful comments on the manuscript.

